# Somatic mutability in cancer predicts the phenotypic relevance of germline mutations

**DOI:** 10.1101/447862

**Authors:** Paolo Provero, Dejan Lazarevic, Davide Cittaro

## Abstract

Genomic sequence mutations in both the germline and somatic cells can be pathogenic. Several authors have observed that often the same genes are involved in cancer when mutated in somatic cells and in genetic diseases when mutated in the germline. Recent advances in high-throughput sequencing techniques have provided us with large databases of both types of mutations, allowing us to investigate this issue in a systematic way. Here we show that high-throughput data about the frequency of somatic mutations in the most common cancers can be used to predict the genes involved in abnormal phenotypes and diseases. The predictive power of somatic mutation patterns is largely independent of that of methods based on germline mutation frequency, so that they can be fruitfully integrated into algorithms for the prioritization of causal variants. Our results confirm the deep relationship between pathogenic mutations in somatic and germline cells, provide new insight into the common origin of cancer and genetic diseases and can be used to improve the identification of new disease genes.

## Introduction

Cancer has been called a disease of the genome since in most cases it is initiated by mutations occurring in somatic cells leading to uncontrolled proliferation and eventually to metastatic invasion of other tissues. On the other hand many diseases, both rare and common, can be caused or favored by mutations in the germline genome. It is thus natural to ask to what extent the same mutations can be associated to cancer and genetic diseases when occurring respectively in somatic or germline cells.

Indeed many cases are known of genes involved in both types of diseases: for example Rasopathies [1] are a family of developmental diseases caused by germline mutations in genes of the Ras/MAPK pathway, which is also recurrently mutated in many cancer types [2]. This observation extends to other cancer related genes, such as ATRX and KDM5C involved in X-linked mental retardation [3], KMT2D in Kabuki syndrome [5], SQSTM1 in Paget disease [6], and DMNT3A in Tatton-Brown-Rahman syndrome [7]. A recent review [8] pointed out the role of mutations of chromatin remodelers in both cancer and neurodevelopmental disorders. However, to our knowledge, the extent to which the mutational spectrum of cancer and genetic disorders overlap has never been investigated in a systematic way.

Recent development in sequencing technologies allow the determination of mutations in patients in a fast and cost-effective way, especially when the sequencing is limited to exons, so that sequencing is now routinely used as a diagnostic and prognostic tool in both genetic diseases and cancer. These development have also allowed the creation of large databases of mutations including many thousands of individuals, providing us with the means to investigate the relationship between somatic and germline pathogenic mutations in a systematic and statistically controlled way.

We thus decided to investigate whether patterns of somatic mutations detected in cancer samples contain information that can be used to predict the involvment of genes in genetic diseases. We chose to tackle the issue in astatistical learning framework, that is to reframe the question as whether it is possible to predict the involvement of a gene in a genetic disease (or more generally an abnormal phenotype due to germline mutations) using the frequency of its somatic mutations in a set of common cancers. In this way we can take advantage of precise statistical methods to accurately quantify the predictive power of the model, and to determine whether such cancer-based predictors can provide new information when combined with more traditional disease-gene prioritization methods, based for example on the frequency spectrum of germline mutations.

## Results

We obtained from the TCGA project [9] the frequency of somatic mutations for 20275 protein-coding genes in 32 cancers. From the Human Phenotype Ontology (HPO) [10] we obtained 355239 association between 1368 phenotypes and 3664 genes (we considered only phenotypes with more than 50 associated genes to avoid problems in fitting logistic models; moreover we limited the analysis to HPO terms classified as *phenotypic abnormality* but not as *neoplasm*).

### The frequency of somatic mutations in cancer predicts the involvment in abnormal phenotypes

To verify whether patterns of somatic mutations in cancer are correlated with involvment in genetic diseases we first built a logistic model in which the regressor variable is the total number of somatic mutations of a gene (TSM), summed over all TCGA samples across all cancer types, and the predicted outcome is the involvment of the gene in the most generic HPO phenotype we consider, namely *phenotypic abnormality* (HP:0000118). Since the length of the coding sequence of a gene correlates with the TSM (r = 0.676) we introduced it as a covariate.

This model turns out to be highly predictive, with an AUC of 0.623 and an ANOVA P-value of 3.64e-102. Importantly TSM is an independent predictor (P = 1.08e-50), while coding sequence length is not (P = 0.606). The odds ratio of the TSM per units of its standard deviation is greater than 1 (OR = 1.47, 95% CI = (1.39 - 1.54), indicating that genes with higher TSM are more likely to be involved in abnormal phenotypes.

### The frequency of somatic mutations in cancer is independent of predictors based on germline mutation frequency

Several disease gene predictors have been developed based on the germline mutation data derived from large exome sequencing projects. To determine whether somatic mutation frequency in cancer is independent of such predictors we fitted a multivariate logistic model which included as regressors, besides TSM and coding sequence length, a predictor derived from germline mutation frequency and one derived from denovo mutation frequency. Multivariate regression allows to determine the significance of the association between each regressor and the outcome, all other regressors being fixed, and thus whether each regressor provides independent predictive power. Specifically, we used the probability of being tolerant of both heterozygous and homozygous loss-of-function variants computed from ExAC [11] and the mutability index based on denovo mutation frequency [12].

As expected this multivariate logistic model is highly predictive of involvment in phenotypic abnormality (AUC = 0.666, ANOVA P = 3.69e-201). Figure 1A shows the odds ratio and P-value associated to each predictor (per unit of its standard deviation). Therefore all three quantities based on cancer, germline and denovo mutation frequencies are independent predictors. Note that the OR value for the mutability index is greater than one, in agreement with the somewhat surprising observation [12] that disease genes are *more* prone to denovo mutations than other genes. Obviously the OR for pNull is <1, since it indicates the probabilty of being *tolerant* of germline mutations.

### Frequency of somatic mutations in cancer correlates positively with embryonic lethality but negatively with cellular essentiality

We used the same model to predict embryonic lethality, using mouse data [13], and essentiality in human cell lines derived from towo large-scale studies [14,15]. For embryonic lethality the results are in line with those found for abnormal phenotypes, as shown in Figure 1B, except that pNull is the strongest predictor, in line with the fact that pNull was created specifically as a predictor of essentiality. Note however that both mutability and TSM do independently contribute to the prediction of experimentally determined essential genes.

**Figure 1:**
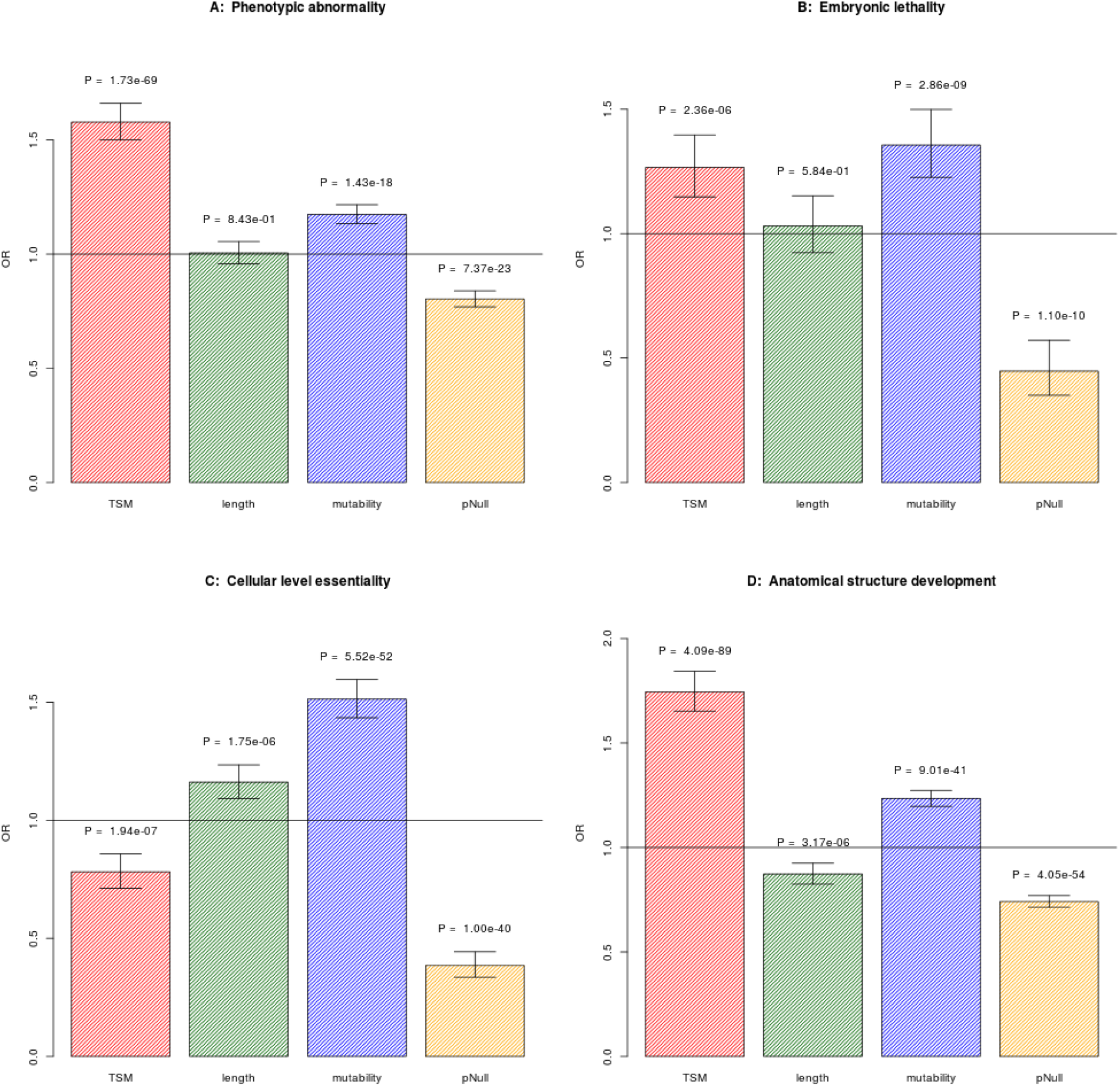
Multivariate logistic models predicting various gene annotations. We show the odds ratio and its 95% confidence level with the corresponding P-value for the four covariates: TSM - total somatic mutations in cancer; length: length of the coding region; mutatbility: mutability index obtained from denovo mutation frequencies; pNull: probability of being tolerant of bot heterozygous and homozygous variants. A: involvment in a generic phenotypic abnormality (HP:0000118); B: Embryonic lethality; C: Cellular level essentiality; D: Involvment in anatomical structure development (GO:0048856)

Regarding essentiality in human cell lines, Figure1C shows the result of fitting the model to the genes that are found essential in both [14] and [15]. Contrary to what happens for abnormal phenotypes and embryonic lethality TSM *negatively* correlates with essentiality. This is telling us that all other predictors being equal, a gene that is highly mutated in cancer is unlikely to be essential at the cellular level, as it is expected since mutations in genes that are essential for cell viability are likely to be negatively selected in tumors.

### The frequency of somatic mutations in cancer predicts developmental gene function

Since most abnormal phenotypes are the result of defects in the execution of developmental programs we hypothesized that TSM should positively correlate with development-related functional annotation. We considered the 191 Gene Ontology terms belonging to the Biological Process ontology and containing the string “development”, and used the same multivariate logistic model to predict such annotation. The contribution of the TSM to the prediction was significant (P < 0.05) in 160 out of 191 GO terms, and in all these cases the OR was greater than 1. Table 1 shows the top 10 terms by P-value associated to the TSM, while complete results are in Supplementary Table 1. Figure 1D shows the coefficients and P-values of all the covariates for “anatomical structure development” (GO:0048856).

**Table 1:** Odds ratio, its 95% confidence interval and P-value associated to TSM for the development-related GO terms (shown are the top 10 terms by TSM P-value).

| Term | OR | CI | P |
| --- | --- | --- | --- |
| anatomical structure development | 1.745 | 1.652 - 1.843 | 4.09e-89 |
| developmental process | 1.700 | 1.614 - 1.792 | 2.21e-88 |
| multicellular organism development | 1.747 | 1.652 - 1.848 | 4.38e-85 |
| system development | 1.697 | 1.604 - 1.797 | 1.61e-74 |
| cellular developmental process | 1.502 | 1.423 - 1.586 | 1.80e-49 |
| nervous system development | 1.639 | 1.533 - 1.752 | 1.99e-47 |
| animal organ development | 1.498 | 1.412 - 1.590 | 7.91e-41 |
| cell development | 1.460 | 1.372 - 1.554 | 1.66e-32 |
| regulation of developmental process | 1.376 | 1.292 - 1.465 | 1.89e-23 |
| neuron development | 1.489 | 1.373 - 1.614 | 7.57e-22 |

### Results for specific phenotypes and tumor types

We then asked for which tumor types and which abnormal phenotypes show the strongest predicitve power of TSM. For each of the 1368 abnormal phenoytpes and each of the 32 tumor types (listed in Supplementary Table 2) we fitted a logistic model with the same regressors as before, except that the TSM is computed only on samples of the specific tumor type.

In 20556 out of 43776 models the TSM term was signifcant (Benjamini-Hochberg FDR < 0.05 computed over all 43776 models). Figure 2 represents the number of phenotypes for which the TSM of each tumor type is a significant predictor (left panel) or the best predictor (right panel) compared to the other regressors, as a function of the mean number of mutated genes per patient in each tumor. The positive correlation result is compatible with a scenario in which the *somatic* mutability of a gene is predictive of involvment in abnormal phenotypes, similarly to what shown [12] for *de novo* germline mutability. Tumors with high mutation rates provide a more accurate sampling of somatic mutability and thus a better predictor of phenotypic effects. We will return to this point in the Discussion. Figure 3 shows the coefficient (natural log of the odds ratio) of the TSM in predicting the most general abnormal phenotypes.

**Figure 2:**
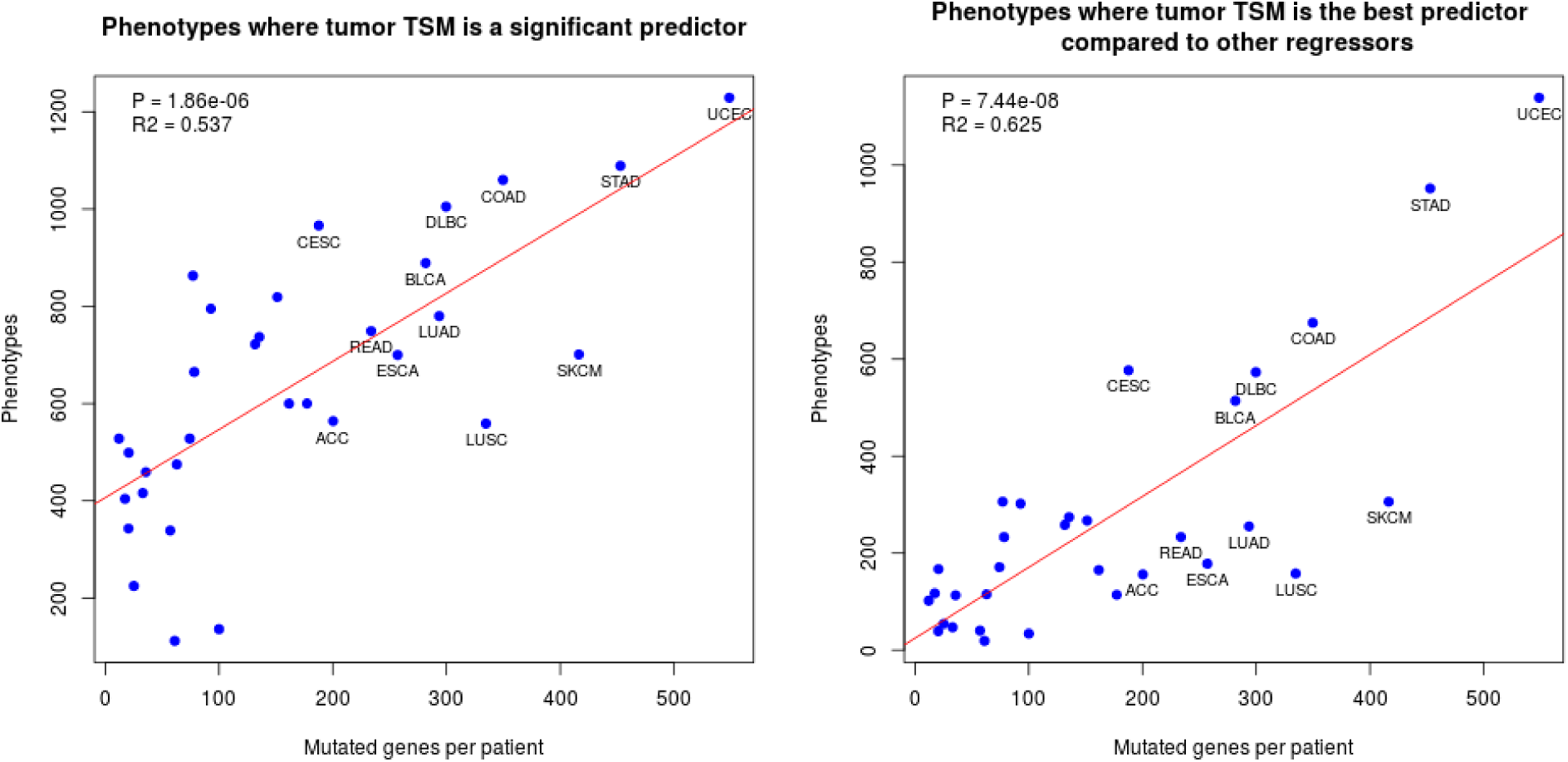
Left: number of HPO phenotypes for which TSM of each tumor type is a significant predictor, independent of the other covariates, as a function of the mean number of mutated genes per tumor sample. Right: number of HPO phenotypes for which TSM is the most significant predictor compared to the other covariates. The P-value and R^2^ are derived from a linear model, with the regression line shown in red. For readability only the tumor types with the highest mutation rates are labelled.

**Figure 3:**
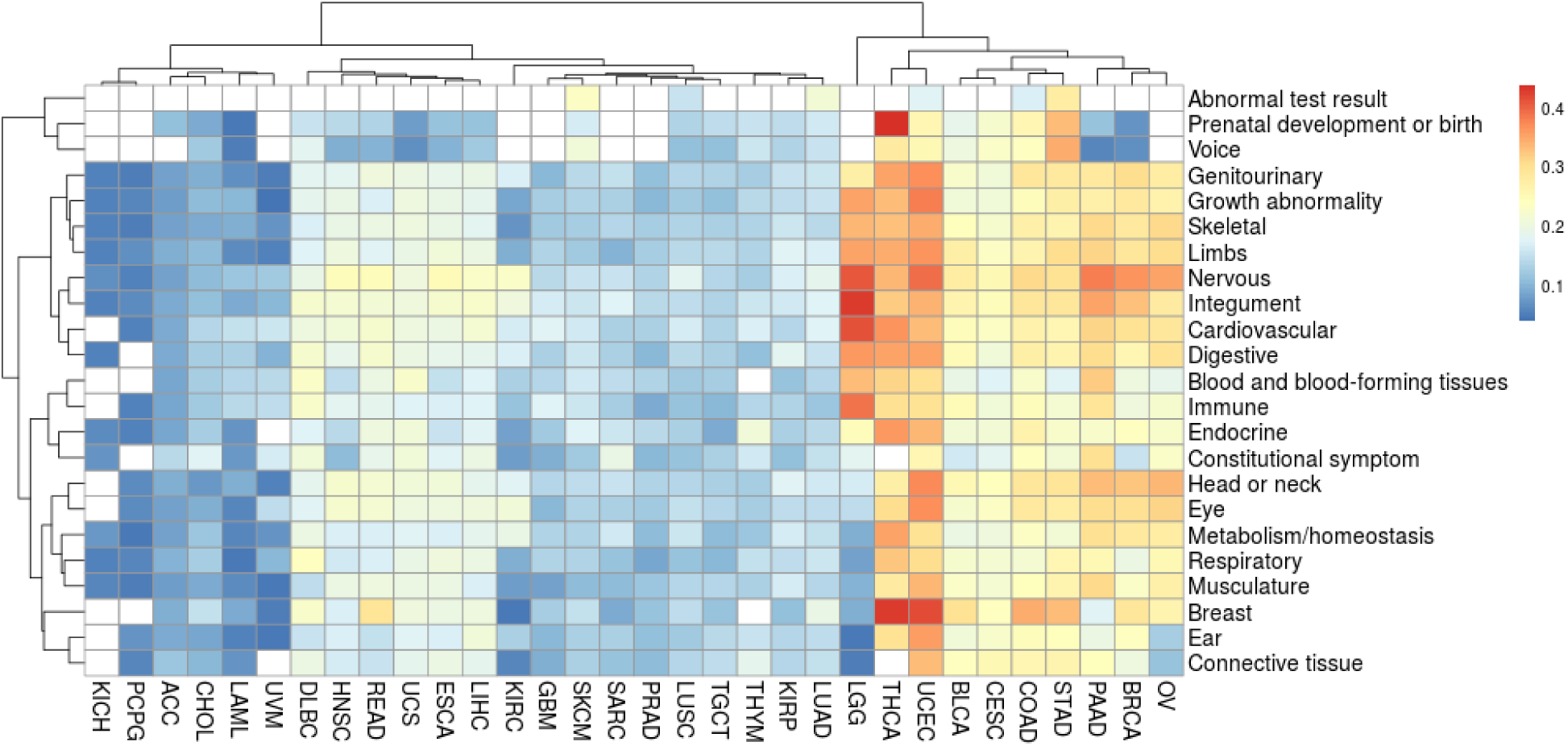
Coefficient of the TSM as a predictor of gene involvment in the most general abnormal phenotypes collected in the Human Phenotypes Ontology (children of the term “phenotypic abnormality”). White cells correspond to cases in which the TSM is not a significant predictor of gene involvement.

### Integrated models for gene prioritization

The previous results demonstrate that patterns of somatic mutations in cancer can be successfully integrated with previously derived indicators derived from germline mutation frequencies to improve the prediction of genes involved in abnormal phenotypes. Specifically we propose to select, for each phenotype, the multivariate logistic model corresponding to the tumor type that gives the highest AUC. The number of phenotypes for which each tumor type is thus selected is shown in the Figure 4.

**Figure 4:**
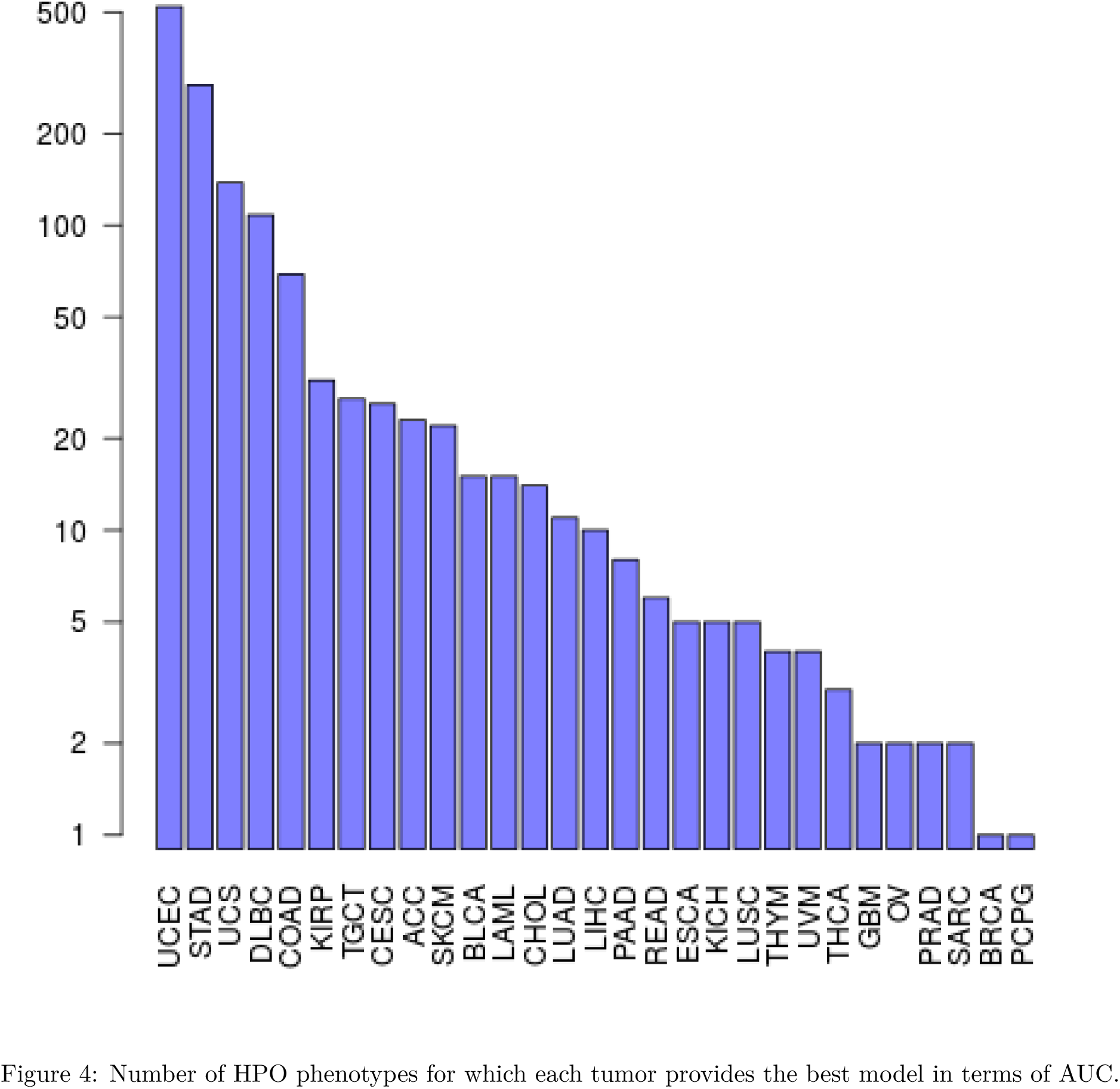
Number of HPO phenotypes for which each tumor provides the best model in terms of AUC.

Table 2 shows the AUC of the final model and the ANOVA P-value for the most general phenotypes, while Table 3 shows the phenotypes with the best AUC, together with the selected tumor type. AUC and P-values for all phenotypes examined are reported in Supplementary Table 3.

**Table 2:** Most predictive model, AUC and ANOVA P-value for the most general abnormal phenotypes (children of phenotypic abnormality - HP:0000118)

| term | best | AUC | P |
| --- | --- | --- | --- |
| Abnormality of the voice | STAD | 0.704 | 1.92e-26 |
| Abnormality of limbs | UCEC | 0.689 | 4.69e-122 |
| Abnormality of the breast | UCEC | 0.683 | 1.01e-24 |
| Abnormality of connective tissue | UCEC | 0.679 | 1.71e-83 |
| Abnormality of head or neck | UCEC | 0.676 | 1.94e-158 |
| Abnormality of the musculature | UCEC | 0.674 | 1.15e-139 |
| Abnormality of prenatal development or birth | STAD | 0.674 | 1.08e-29 |
| Abnormality of the nervous system | UCEC | 0.673 | 2.73e-181 |
| Abnormal test result | SKCM | 0.673 | 9.65e-04 |
| Abnormality of the skeletal system | UCEC | 0.672 | 1.62e-146 |
| Abnormality of the ear | UCEC | 0.671 | 9.00e-105 |
| Abnormality of the eye | UCEC | 0.670 | 4.34e-142 |
| Abnormality of the genitourinary system | UCEC | 0.658 | 1.37e-92 |
| Abnormality of the digestive system | UCEC | 0.658 | 2.76e-106 |
| Growth abnormality | UCEC | 0.656 | 2.72e-100 |
| Abnormality of the cardiovascular system | UCEC | 0.656 | 2.08e-101 |
| Abnormality of the respiratory system | UCEC | 0.655 | 7.41e-74 |
| Abnormality of the immune system | PAAD | 0.655 | 1.03e-65 |
| Abnormality of the integument | UCEC | 0.654 | 7.50e-99 |
| Abnormality of blood and blood-forming tissues | DLBC | 0.648 | 2.71e-46 |
| Abnormality of the endocrine system | UCEC | 0.646 | 8.04e-46 |
| Constitutional symptom | PAAD | 0.642 | 1.08e-25 |
| Abnormality of metabolism/homeostasis | UCEC | 0.629 | 5.60e-62 |

**Table 3:** AUC and ANOVA P-value for the 10 phenotypes with the highest AUC

| term | best | AUC | P |
| --- | --- | --- | --- |
| Thick eyebrow | STAD | 0.813 | 3.80e-18 |
| Skin tags | STAD | 0.807 | 1.09e-15 |
| Cutis marmorata | READ | 0.805 | 5.71e-11 |
| Narrow face | UCEC | 0.804 | 4.50e-12 |
| Mitral valve prolapse | STAD | 0.802 | 2.15e-12 |
| Congenital hip dislocation | TGCT | 0.801 | 3.01e-10 |
| Absent toe | STAD | 0.800 | 4.31e-11 |
| Broad thumb | STAD | 0.797 | 9.87e-12 |
| Dental malocclusion | CESC | 0.796 | 1.92e-14 |
| Long eyelashes | STAD | 0.794 | 1.55e-12 |

### Combining phenotypes to predict disease genes

We asked how our predictor would perform when applied to genes associated to genetic diseases, rather than individual phenotypes. The HPO site provides a table associating a list of phenotypes to ORPHANET [16] and OMIM [17] entries. A disease gene predictor can thus be obtained by suitably combining the predictions for its associated phenotypes. We chose to aggregate the phenotype predictors by rank products: the score of a gene as a candidate for a disease is the geometric mean of the ranks of the gene as a predictor of the phenotypes associated to the disease. Distribution of combined ranks for true gene/disease associations are significantly higher than random associations when combined score is lower than 0.1, as shown in Figure5; we obtained similar results for OMIM entries (Supplementary Figure 1).

**Figure 5:**
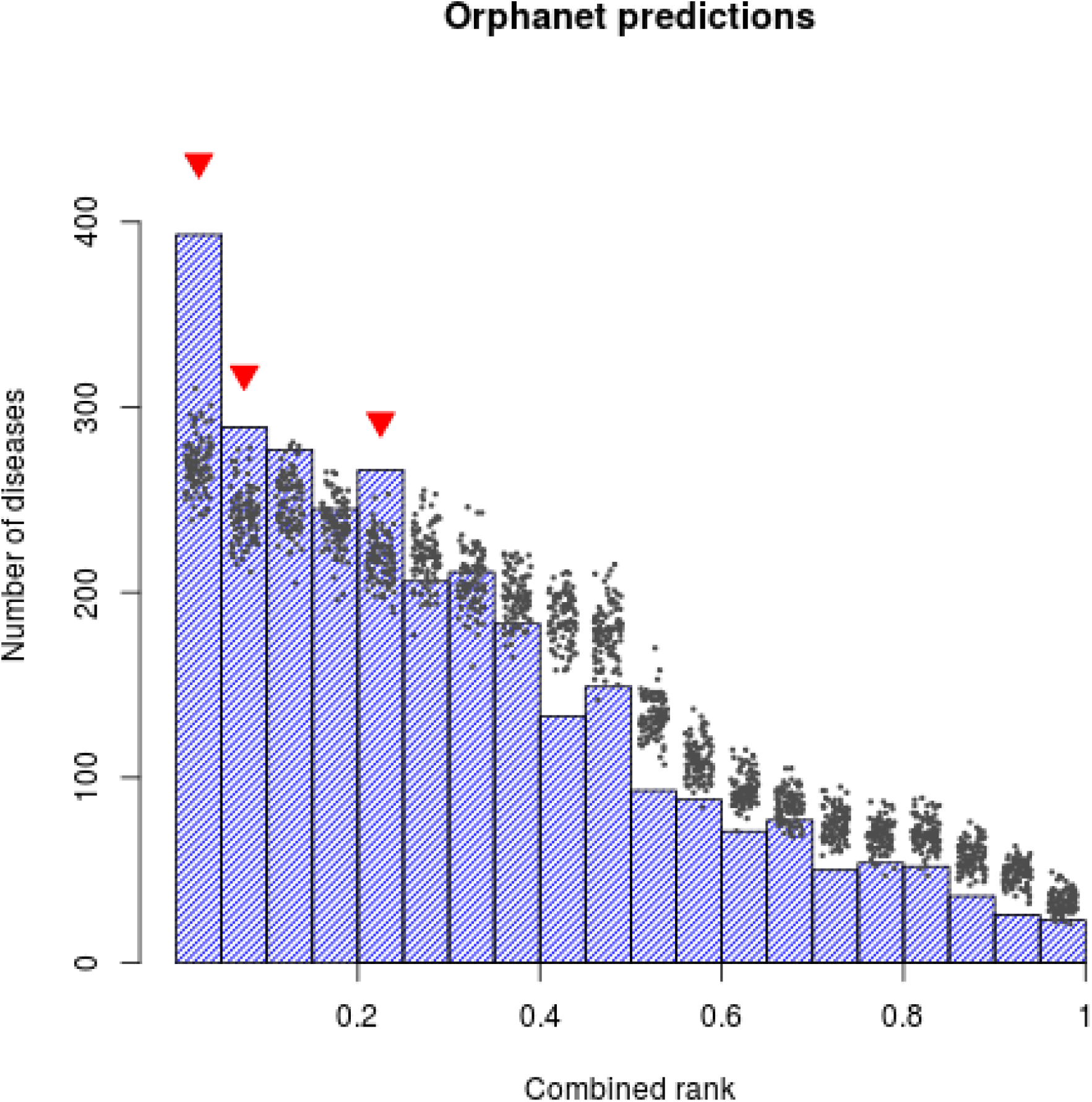
Distribution of combined ranks for orphanet disease-gene associations. Blue bars indicate the actual value of combined rank for every disease-gene association reported in the Orphanet database. Black dots represent the realization of combined scores of 100 randomizations of gene-disease associations. Red triangles mark the bins for which the actual combined rank is lower (better) than the corresponding randomizations (z score test, p < 0.01)

In agreement with the central role of disrupted developmental processes in generating abnormal phenotype we observed that the age of onset of disease influences the distribution of combined ranks for the relative disease genes; in particular, lower combined ranks are associated with early-onset diseases, whereas we observe highest, hence less predictive, ranks for diseases arising during adolescence (Kruskal-Wallis P = 0.01; Supplementary Fig. 2). An interactive application disease-gene prediction is available at https://dawe.github.io/discan/ as jupyter notebook.

## Discussion

We have shown that the large-scale assessment of somatic mutations in tumors is not only useful in understanding the genetics of cancer, but also in an unexpected context, namely in predicting *germline* mutations responsible for phenotypes and diseases. Genes that are recurrently mutated in tumors are more likely to be involved in abnormal phenotypes when mutated in the germline. This connection had been suggested based on specific cases, but here we have shown that it is generally true in a statistically controlled way. Notably, the link between disease-causing germline mutations and somatic mutations in cancer was recently exploited in [18] in the opposite direction, namely to predict driver somatic mutations based on their pathogenic role in Mendelian diseases.

These results have practical implications and raise conceptual issues. From the practical point of view, they suggest that TSM profiles could be profitably integrated with other sources of information (germline variation patterns, functional annotation, biomolecular networks, etc.) in developing tools for the prioritization of disease genes.

Conceptually, the emerging picture is that mutability is a strong predictor of involvment in abnormal phenotypes. Genes involved in a phenotypic abnormality must have two features: they must be highly mutable [12], but their mutation must not affect cell-level viability. But the frequency of somatic mutations in cancer depends precisely on these two features: a gene will be frequently mutated in tumors if it is easily mutated and its mutation does not kill the cell. Therefore human tumors provide us with a large scale natural experiment in which the two hallmarks of disease genes are probed simultaneously. Indeed the tumors withthe highest predictive power are the one with the greater number of mutations, since these tumors are able probe more deeply the mutability spectrum of human genes.

Importantly the correlation between mutability and disease genes does not suggest that gene function is not relevant: on the contrary, mutability, both *de novo* in the germline and somatic in tumors, is a strong predictor of function. In particular highly mutable genes are often involved in developmental programs whose disruption leads to inherited abnormal phenotypes and diseases.

Our analysis was performed at the gene level, in particular because the variables we use together with the TSM in bulding our predictors are defined at such level. It would be interesting to investigate at the single base level whether the same specific mutations that appear in cancer as somatic are also causative of phenotypic abnormalities.

## Methods

### Data

#### Somatic mutations in cancer

Mutation data for 20275 genes and 32 tumor types were collected from TCGA using the RTCGAToolbox Bioconductor package. From these we derived for each gene and tumor type:

- the number of patients with a mutation in their primary tumor
- the same restricted to non-silent mutations
- the same restricted to silent mutations

Only variants classified as *Frame*_*Shift*_*Del*, *Frame*_*Shift*_*Ins*, *In*_*Frame*_*Del*, *In*_*Frame*_*Ins*, *Missense*_*Mutation*, *Nonsense*_*Mutation*, *Silent*, *Splice*_*Site* and *Translation*_*Start*_*Site* were considered, implying that these are all in the coding region (or within 2bps of an exon in the case of splice site variants). The bivariate model predicting involvment in abnormal phenotype was performed considering all mutations, only non-silent or only silent mutations. The model using all mutations was slightly more predictive than the one using non-silent mutations only, and both were more predictive than the one using silent mutations only. Therefore in all following models we used the frequency of all mutations.

#### Coding sequence length

Coding sequence length of all human genes was defined as the sum of the lengths of all annotated exons assigned to the gene, irrespective of alternative isoforms. Exon annotation for human genome version hg19 was obtained from the “TxDb.Hsapiens.UCSC.hg19.knownGene” Bioconductor package.

#### Genes involved in abnormal phenotypes and in developmental processes

Gene-phenotype associations were downloaded from the Human Phenotype Ontology web site together with the ontology structure. Gene associations were expanded to include all ancestors of the term directly associated to each gene. We thus obtained 431598 associations involving 7848 phenotypes and 3682 genes. To prevent overfitting we built models only for phenotypes with at least 50 associated genes (see also the subsection on cross-validation below).

We considered only the “phenotypic abnormality” (HP:0000118) HPO annotation and all its dsecendants in the HPO graph. We also excluded the phenotype “Neoplasm” (HP:0002664) and all its descendants since the association between genes somatically mutated in cancer and genes involved in cancer predisposition is expected and might be explained by different mechanisms. All these criteria led to a set of 1368 HPO phenotypes on which the logistic models were fitted.

Genes annotated to developmental processes were obtained from the Gene Ontology annotation reported in the “org.Hs.eg.db” Bioconductor package. We considered the GO terms in the Biological process ontology matching the string “development”.

#### Other mutability measures

The mutability index (MI) for each gene was derived from Suppl. Table 3 of [12] as the average MI of all the exons of a gene weigthed by exon length. The probability of being tolerant of both heterozygous and homozygous lof variants (pNULL) was obtained from Supplementary Table 13 of [11].

#### Disease genes

Annotation of ORPHANET and OMIM diseases and genes were downloaded from Orphanet portal ([16], latest access 27/01/2018). Since our analysis is limited to a subset of HPO terms, annotated phenotypes may be excluded. For this reason, we projected annotated HPO terms to the first element in the shortest path from an annotation to the HP:0000118 term (“phenotypic abnormality”) which is included in our set. Once a set of uniquely projected terms is obtained, the combined score R of a gene for a disease is calculated as

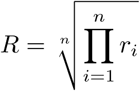

where *r_i_* is the rank of a gene for the i-th phenotype and n is the number of phenotypes under consideration. Random scores are calculated by shuffling gene names and diseases in the gene/disease associations.

### Statistical analysis

#### Logistic regression

The predictive power of the regressors was evaluated using standard multivariate logistic regression. All regressors (total somatic mutations, cds length, mutability index, pNULL) were standardized so that the reported odds ratio refer to one unit of standard deviation. Missing values were imputed using KNN imputation as implemented in the “bnstruct” CRAN package. AUCs were evaluated with the “ROCR” package.

#### Cross-validation

Overfitting of the multivariate logistic model is unlikely since the number of genes associated to the phenotypes we consider is at least 50 and we have at most 4 regressors. To confirm this we computed cross-validated AUCs using the cvAUC package and 5-fold cross-validation. For 1358 out of 1368 cases the AUCs computed on the training set were inside the 95% confidence interval determined by cross-validation, confirming that overfitting is not a relevant issue.

## Supplementary material

- **Supplementary Table 1**. Multivariate logistic models predicting the annotation to Gene Ontology Biological Process term related to development. For each of the four regresstors described in the text we report the odds-ratio (OR) in units of the standard deviation, its 95% confidence interval (OR_ci_high, OR_ci_low) and its associated ANOVA P-value (P).
- **Supplementary Table 2**. Tumor types analyzed with their abbreviations.
- **Supplementary Table 3**. Best predictive model and its performance for all HPO phenotypes analyzed. The P-value is computed with ANOVA by comparing the full model to the one with intercept only.
- **Supplementary Figure 1**. Distribution of combined ranks for OMIM disease-gene associations, as in Fig.5
- **Supplementary Figure 2**. Distribution of combined rank in according to the Age of Onset of disease. The boxplot represents the distribution of combined ranks of disease-gene associations from Orphanet database, stratified by Age of Onset. Diseases without a reported AoO were excluded from the graph. Kruskal-Wallis test shows that the correlation between age of onset and combined rank is significant (P = 0.01)

## Acknowledgements

This work was supported by the Italian Ministry of Health under the “5 per mille” program

